# Differential modulation on neural activity related to flankers during face processing: a visual crowding study

**DOI:** 10.1101/2023.05.17.541244

**Authors:** Zeguo Qiu, Dihua Wu, Benjamin J. Muehlebach

## Abstract

The modulations of visual awareness on the processing of emotional faces have been mainly investigated in masking experiments where stimuli were presented for different durations as an integral process to the masking technique. In this visual crowding study, we manipulated the perceivability of a central crowded face (either a fearful or a neutral face) by varying the similarity between the central face and the surrounding flanker stimuli. We presented participants with pairs of visual clutters and recorded their electroencephalography during an emotion judgement task. In an upright flanker condition where both the target face and flanker faces were upright faces (high similarity), participants were less likely to report seeing the central target face, and their P300 was weakened, compared to a scrambled flanker condition where scrambled face images were used as flankers (low similarity). Additionally, at around 120ms post-stimulus, a posterior negativity was found for the upright flanker condition, compared to the scrambled flanker condition, however only for fearful face targets. We concluded that early neural responses seem to be affected by the perceptual characteristics of both target and flanker stimuli whereas neural activity at a later stage is associated with post-perceptual evaluation of the stimuli in this visual crowding paradigm.

## Introduction

Emotional expressions such as fear and anger convey a person’s emotional experience. Fearful facial expressions, in particular, often indicate potential threatening events in our surrounds. This is thought to be one of the reasons why fearful faces attract attention and break into awareness more easily than neutral facial expressions (for reviews see [1,2]). Electrophysiological recordings have been a useful tool in examining the neural activity during the processing of emotional faces. Specifically, it has been shown that emotional faces enhance both early and later-stage event-related potentials (ERPs), compared to their neutral counterparts (for a review see [2]), even when unattended [3,4,5]. For example, presenting face images in pairs, we previously showed that a fearful face in the pair increased early posterior ERPs (i.e., N2) compared to a neutral face, both when it was spatially attended and when spatially unattended [4].

More recent studies have shown that the increased attentional attraction caused by fearful faces may be dependent on a variety of factors including task-relevancy [6] and the perceivability of the stimuli [4,6,7,8]. For example, in a backward masking experiment, we presented participants with either nearly invisible (subliminal viewing) or clearly visible (supraliminal viewing) face pairs. We found that the N2-posterior-contralateral (N2pc), an ERP indexing attentional capture, was elicited by a fearful target face only when stimuli were presented supraliminally [6]. Further, when the task required participants to attend to non-facial feature that was superimposed onto the face images, the N2pc for the fearful face was no longer observed. Therefore, we concluded that attentional capture by fearful faces requires visual awareness and is contingent upon task-relevancy of the faces [6]. Similar findings have been replicated since [9,10,11].

A common criticism of the backward masking paradigm in the awareness literature is that stimuli must appear for different durations in the subliminal and supraliminal viewing conditions (e.g., 16ms vs. 266ms in [4]). ERPs time-locked to stimulus onset, especially early ERPs, may be sensitive to perceptual differences between conditions in this case. An alternative way of implementing different conditions of awareness is using a visual crowding paradigm. Visual crowding is a phenomenon where the perception of a target stimulus is impaired when it is closely surrounded by other stimuli, typically referred to as flankers. The perceivability of a central crowded target can be varied by modifying the flankers (e.g., their location or their complexity; [12]). Different from the backward masking paradigm, stimulus durations are kept the same across conditions.

Studies on the effects of visual crowding on face processing are rather limited. In a study by Kalpadakis-Smith and colleagues [13], by varying the levels of target-flanker similarity for the face stimuli, it was found that participants’ identification of a central crowded face was impaired most significantly when flankers were upright faces (i.e., the most similar to the target). When the flankers were inverted faces, the identification of the central target face improved [13]. Another study by Gong and Smart [14] found that when the flankers expressed an emotion (i.e., happy) that was different from the central target face (i.e., angry), participants were worse at judging the emotion of the central face, compared to when the flankers were neutral faces and when the flankers’ emotion matched the target. Based on the above literature, in a visual crowding paradigm, using actual face images presenting an emotional expression that is different from the central target face may be used as flankers to maximise suppression of target face awareness.

The present ERP study aims to examine the modulation of visual awareness on the neural activity for fearful and neutral face processing by using the visual crowding paradigm. Specifically, in our two flanker conditions, upright angry faces and scrambled angry faces were used as flankers, respectively. Upright flankers were expected to impede participants’ awareness of the central target face more strongly than scrambled flankers [13,15]. As we were primarily interested in the neural correlates of awareness when comparing the two flanker conditions, we focused on two time ranges that are associated with visual awareness: the N2 and the P300 time ranges. Specifically, in the N2 time range, stimuli that are consciously processed are usually associated with more negative signals, compared to unconscious stimuli. This negativity for conscious stimuli is termed the visual awareness negativity (VAN) and is suggested to index an early perceptual stage of awareness (for a review please see [16]). Similarly, the P300 has been found to be larger for conscious relative unconscious stimuli [17], which is suggested to reflect a later reflective stage of awareness [16].

In the current study, we expected to find a VAN and an enhanced P300 for central target faces in the scrambled flanker condition, compared to the upright flanker condition. We predicted that a fearful expression of the target faces should further enhance the VAN and the P300, compared to neutral faces, in line with previous findings [4,18].

## Material and methods

### Participants

We determined the sample size based on previous studies on visual crowding. As most of the behavioural studies used small to moderate sample sizes ranging from 4-25 [13,15,19,20], we decided to choose a sample size closer to a previous ERP study (i.e., 18 participants in [21]) and recruited 22 participants at the University of Queensland.

Participants were compensated with course credits. Data from two participants were excluded from analyses due to excessive eye-movements in the data. Thus, 20 participants constituted the final sample (*M*_*age*_ = 23.6 years, *SD*_*age*_ = 4.0 years, 16 females, 4 males).

This study was approved by the University of Queensland ethics committee. All participants provided written and verbal consent prior to the study.

### Apparatus and experimental stimuli

All stimuli were presented on a 19” colour LCD monitor (resolution: 1280 x 1024 pixels) with a viewing distance of 50cm. We monitored participants’ monocular gaze position using the Eyelink 1000 plus system (SR Research Ltd., Canada) with a spatial resolution of <0.01° and a sampling rate of 500 Hz. The program was run in PsychoPy3 [22].

Face images were obtained from the Radboud Faces Database [23]. Fearful, neutral and angry face images from 16 models (8 females, 8 males) were used. Each individual face image was first rendered greyscale and then cropped into oval shapes (3.2° x 4.1° in visual angle; see Figure 1a). We used a scrambled filter tool (http://telegraphics.com.au/sw/product/scramble) to make flanker stimuli for the scrambled flankers condition. Specifically, each cropped angry face image was sliced into 208 small squares and randomly shuffled (Figure 1a). All photo editing was done in Photoshop 2021 version 22.4.0 (Adobe Systems, San Jose, CA).

**Figure 1.**
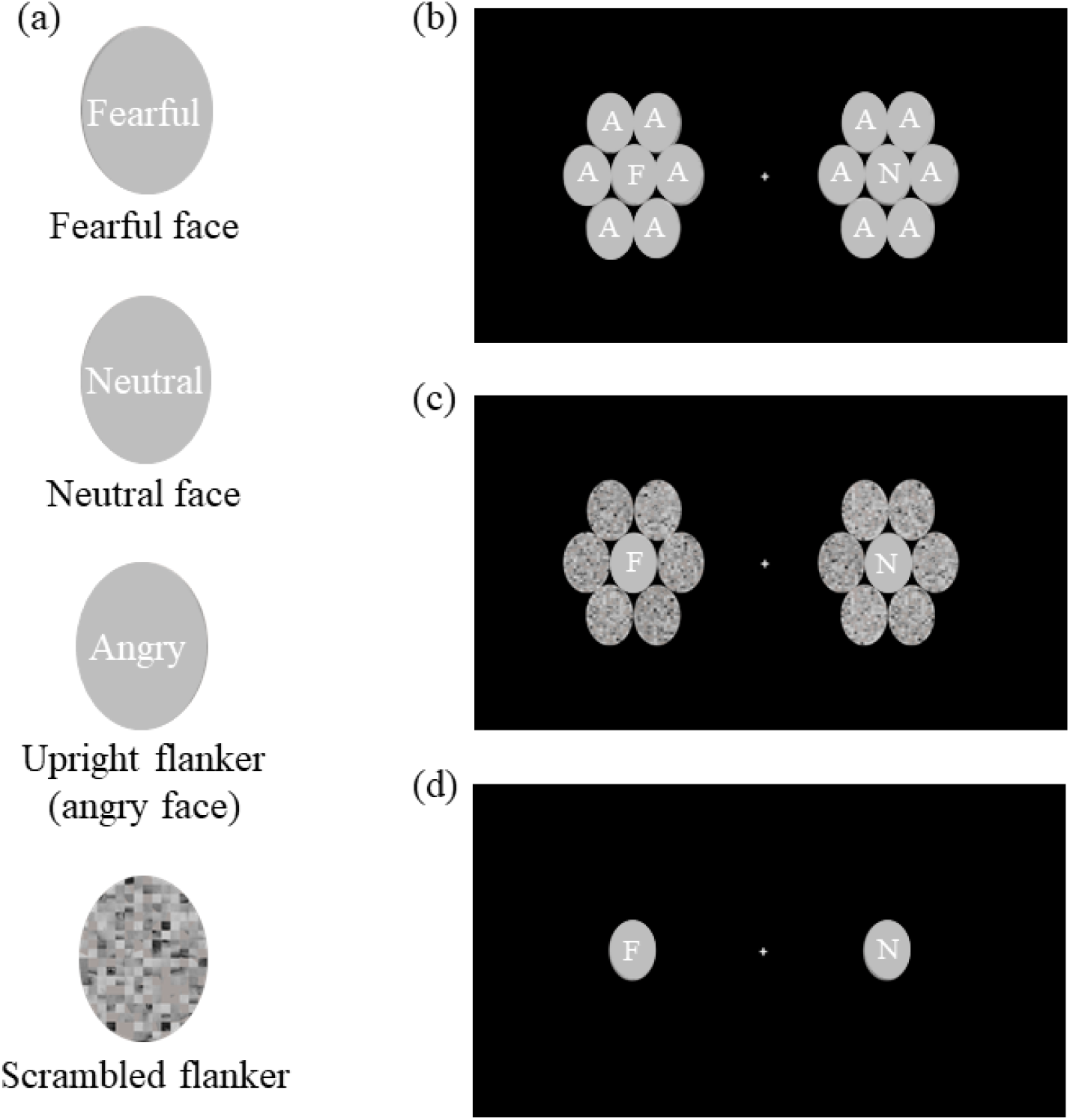
(a) Examples of individual face stimuli. (b) Example stimuli in the upright flanker condition. (c) Example stimuli in the scrambled flanker condition. (d) Example stimuli in the no-flanker condition. Note: The face stimuli are covered and de-identified in this picture in accordance with bioRxiv policies. Actual face images were used in the experiment.

Each target screen consisted of two clutters of face images with one clutter on each side of the screen. In each clutter, there was a central target face that could either be a fearful face or a neutral face, surrounded by six flanker faces. The emotion of the central faces in the two clutters were independently and randomly determined by the program such that the central face (a fearful or a neutral face) on the left clutter could be paired with a fearful or a neutral face in the right clutter equiprobably. In the upright flanker condition, six angry faces from different models were used as flankers (see Figure 1b). In the scrambled flanker condition, six scrambled faces from different models were used as flankers (see Figure 1c). In the no-flanker condition, no flankers were presented (see Figure 1d).

The locations (visual angles) of the central target faces were determined in a pre-experiment calibration process for each participant (please see Procedure below), and they ranged from 7° to 14° (with reference to the screen centre) across participants. The locations of the six flanker stimuli were, with reference to the location of the central target faces, as follows: (target face visual angles ±3.2°, 0) or (target face visual angles ±1.6°, ±3.6°). As can be seen in Figure 1b & 1c, there was no obvious gap between the central target faces and the flankers, consistent with previous studies [13,15].

### Procedure

The study consisted of two parts: a pre-experiment calibration process and the experiment. Before both the calibration and the experiment, the eye tracker was calibrated with a 5-point calibration.

#### Calibration process

Because it has been repeatedly shown that visual crowding effects vary across individuals, we determined the visual angles for the visual stimuli individual-by-individual during a pre-experiment calibration process [24]. Prior to the experiment, we asked the participants to complete a calibration step to determine the visual angles or the locations of the central target faces. Faces were always presented in pairs with one single face on each side of the screen (same as the no-flanker condition used in the official experiment; Figure 1d). In each trial, participants were asked to fixate at the screen centre and judge whether a face presented on the attended side of the screen (counterbalanced across participants) was a fearful or neutral face. In order to make sure that participants fixated at the screen centre and did not directly look at the target face, we used the eye-tracker to monitor participants’ gaze: when participants’ gaze was detected to have moved away from the central fixation point (with a region limit of 5° x 5°), the target face would be replaced with a scrambled image.

Task responses were made by using the Up and Down arrow keys with participants’ right hand. The mapping between the keys and the responses (fearful or neutral) was counterbalanced across participants.

At the beginning, all faces were presented at ±17°, and participants completed blocks of 20 trials. At the end of each block, if the average accuracy of the block was below 90%, the visual angles of the faces were decreased by 1° (i.e., face images were presented closer to the screen centre) and participants were asked to complete another block. Once the average accuracy in a block was 90%, we terminated the calibration process and used the visual angles from that block in the subsequent official experiment.

#### Experiment

After the calibration process, the official experiment began. For the first half of the experiment, participants were instructed to attend to either the left or the right screen side, which was randomly determined by the experiment program. For the second half of the experiment, they attended to the other screen side. Each trial started with a fixation screen for a randomly selected duration between 500 and 800ms, followed by the target screen presented for 500ms (Figure 2). Participants were asked to judge the emotion of the central target face on the attended screen side, while keeping fixated at a fixation point at the screen centre. Similar to the calibration process, when the faces were presented on the screen, if participants’ gaze was detected to have moved outside the central fixation region (5° x 5°), the target face was immediately replaced with a scrambled face image. As a result, participants never saw the target face with foveal vision, which is a crucial requirement for visual crowding effect to occur. The trials where participants’ eyes were not fixated at the screen centre were skipped, and no response was recorded. For trials where eyes were fixated at the screen centre, the target screen was followed by a blank screen of 100ms, after which participants were asked to use the Up and Down keys with their right hand to indicate the emotion of the target face. The response mapping was identical to that in the calibration process and counterbalanced across participants. Then, participants needed to use the Q and W keys with their left hand to indicate whether they have seen the target face in that trial (Q key = *seen*, W key = *unseen*). This question acted as a subjective report of awareness. A blank screen of 1000ms was presented after a response was made, which terminated the trial.

**Figure 2.**
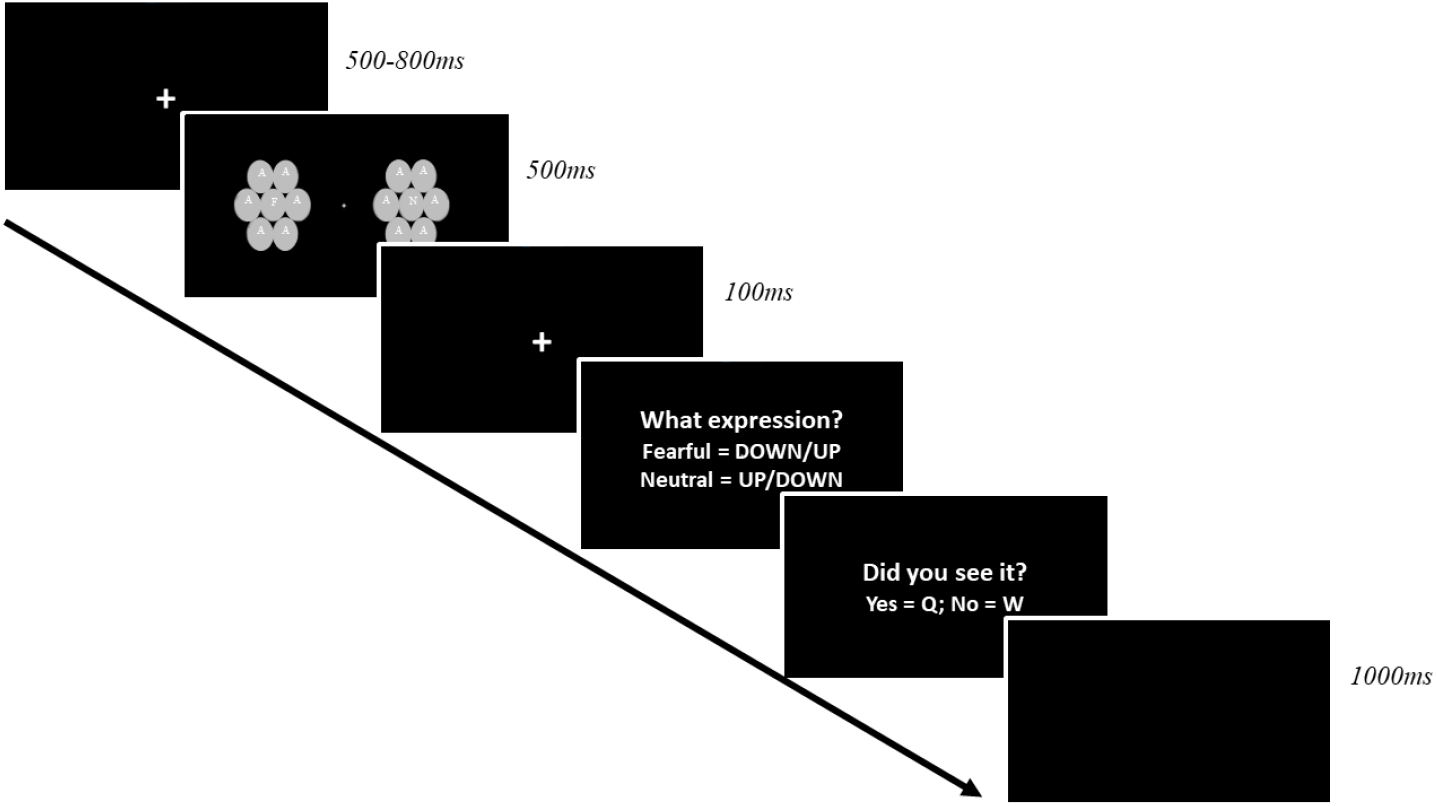
The full procedure of a trial.

There were in total ten blocks with 96 trials in each block. There were 320 trials in each of the three flanker conditions. Breaks were allowed between blocks.

### EEG recording and data pre-processing

Continuous EEG was acquired at 500 Hz using the BrainProducts 32-channel system (Brain Products, Germany) using the international 10–20 configuration. During recording, EEG signals were band-pass filtered between 0.01-40 Hz, and a notch filter of 50 Hz was used to remove power line noise. Recordings were referenced online to a reference electrode taped to participants’ left ear. Impedances were kept below 20kΩ.

Pre-processing of the EEG data was performed with EEGLAB [25] and ERPLAB [26]. We interpolated broken electrodes and electrodes that produced sustained noise throughout the experiment. Signals were filtered from 0.1 to 30 Hz, notch-filtered at 50 Hz to remove line noise and then re-referenced to the average of all electrodes. EEG were segmented into epochs with a time window of 600ms from the onset of the target screen, baseline corrected using the 100ms pre-stimulus interval. As mentioned above, epochs where participants broke fixation were automatically removed. We further visually inspected the epochs on a trial-by-trial basis and removed epochs that contained eye blinks and muscle-related artefacts. Two participants were excluded due to excessive eye-movement in their data. All remaining participants had more than 40 trials in every condition of interest. On average, 73% epochs were kept across participants.

### Data analysis

#### Behavioural responses

For the emotion judgement task, we calculated the accuracy (percent correct) for each flanker and target emotion condition, and performed a 2(emotion: fearful, neutral) x 3(flanker: upright, scrambled, no-flanker) repeated-measures ANOVA. We also applied the signal detection theory method and calculated the d-prime (d’) and criterion (c) for all participants, using fearful face as target signal and neutral face as noise. The d’ ranges from 0 to usually 2 and above, with a larger value indicating a better discriminability of the target. In our case, a negative c would indicate a response bias towards a “fearful face” response whereas a positive c would indicate a response bias to the “neutral face” response.

For the subjective report of awareness, we calculated percentage of giving a seen response for each flanker and target emotion condition, and also performed a 2(emotion) x 3(flanker) repeated-measures ANOVA.

#### Mass univariate analysis

We analysed the ERPs using the factorial mass univariate toolbox [27] and the mass univariate toolbox [28]. Because there were large perceptual differences between the no-flanker condition and the other two flanker conditions, i.e., the total number of stimuli, we excluded the trials from the no-flanker condition, and only examined differences between the two flanker conditions in ERP analyses. Therefore, we conducted a 2(target emotion: fearful, neutral) x 2(unattended emotion: fearful, neutral) x 2(flanker: scrambled, upright) ANOVA. The unattended emotion was included as a variable to examine whether the emotion of the central face in the unattended clutter affected the ERPs, in line with our previous study [4].

For significance testing, we used the cluster-based permutation tests to correct for multiple comparisons (2000 permutations). The threshold for cluster formation was set at an alpha level of 0.05, and electrodes were considered as spatial neighbours if they were within ∼5cm of each other. The mean number of spatial neighbours was 2.5. For statistical significance, we used a family-wise alpha level of 0.05. Follow-up cluster-based permutation two-tailed *t*-tests were conducted where necessary.

## Results

### Behavioural results

#### Emotion judgement task

A 2(target emotion) x 3(flanker) ANOVA on task accuracy revealed a significant main effect of flanker, *F*(2, 38) = 11.41, *p* < .001, *n*_*p*_^*2*^ = 0.38, such that both the upright flanker (*M* = 0.73, *SD* = 0.11) and the scrambled flanker (*M* = 0.74, *SD* = 0.10) conditions were associated with a significantly lower accuracy, compared to the no-flanker condition (*M* = 0.80, *SD* = 0.07), *ps* = .004 (Bonferroni adjusted). There was no difference between the upright flanker and scrambled flanker conditions, *p* = 1.

The interaction between flanker and target emotion was also significant, *F*(2, 38) = 13.98, *p* < .001, *n*_*p*_^*2*^ = 0.42. Follow-up *t*-tests using Bonferroni correction for multiple comparisons showed that, only in the scrambled flanker condition, the task accuracy was much lower when the target face was a fearful face (*M* = 0.65, *SD* = 0.17), compared to when it was a neutral face (*M* = 0.83, *SD* = 0.12), *p* = .002. The lower accuracy for fearful face targets could be driven by a response bias towards indicating a neutral face in the scrambled flanker condition. No significant differences were found in other comparisons, *ps* > .831.

To further investigate whether there was a response bias in the behavioural data, we applied the signal detection theory to the data and calculated d’ and c separately for the three flanker conditions, treating fearful face as the target. Results showed that the d’ was the highest in the no-flanker condition (1.31), followed by the scrambled flanker condition (1.09), and it was the lowest in the upright flanker condition (0.96). These results indicated that participants were best at discriminating a fearful face from a neutral face in the no-flanker condition and the worst at the task in the upright flanker condition. Additionally, we found that the c was close to 0 in both the no-flanker condition (0.01) and the upright flanker condition (-0.02), but it was much higher than 0 in the scrambled flanker condition (0.31).

Therefore, it seems that participants had a tendency to report “neutral face” in the scrambled flanker condition, which could explain the higher accuracy for neutral faces in this condition, as reported above.

#### Subjective report of awareness

We ran a 2(target emotion) x 3(flanker) ANOVA on the percentage of giving a seen response and found a significant main effect of flanker, *F*(2, 38) = 6.83, *p* = .008, *n*_*p*_^*2*^ = 0.27, whereby participants were the least likely to give a “seen” response in the upright flanker condition (*M* = 0.68, *SD* = 0.29) than any other condition, *ps* < .022 (Bonferroni adjusted). There was no difference between the no-flanker condition (*M* = 0.85, *SD* = 0.10) and the scrambled flanker condition (*M* = 0.78, *SD* = 0.22), *p* = .533. The main effect of emotion was also significant, *F*(1, 19) = 10.92, *p* = .004, *n*_*p*_^*2*^ = 0.37, such that a fearful face (*M* = 0.79, *SD* = 0.19) was associated with a higher likelihood of being seen, compared to a neutral face (*M* = 0.75, *SD* = 0.19).

The interaction between target emotion and flanker was significant, *F*(2, 38) = 12.15, *p* < .001, *n*_*p*_^*2*^ = 0.39. Follow-up comparisons showed that in both the no-flanker condition and the upright flanker condition, fearful faces were associated with a higher likelihood of being seen, compared to neutral faces, *ps* = .001 (Bonferroni adjusted). However, the difference between emotion was not significant in the scrambled flanker condition, *p* = .710 (Bonferroni adjusted). Therefore, it seems that fearful face targets were more likely to be seen, compared to neutral face targets, when they were presented on its own (without flankers) and when they were presented with upright angry faces.

### Mass univariate analysis results (ERPs)

ERPs were analysed under the mass univariate analysis framework. A 2(target emotion) x 2(unattended emotion) x 2(flanker) revealed a significant main effect of flanker, and a significant three-way interaction, *Fs* > 4.38, *ps* < .030 (Figure 3a & 3b). Specifically, signals in the scrambled flanker condition were more positive than signals in the upright flanker condition over a posterior cluster (C4, CP5/6, P3/4, PO3/4, PO7/8, O1/2, Oz) between 370 and 594ms (temporal peak: 470ms at electrode P4, *F* = 106.30, *p* < .001), see Figure 3a & 3c. Simultaneously, we found a negativity towards the scrambled flanker condition at frontal and temporal electrodes (F3/4, F7/8, FC5/6, T7/8). The posterior positivity over several parietal electrodes for the scrambled flanker condition may reflect an enhanced P300 for this condition, compared to the upright flanker condition where awareness of the target face was impeded more strongly. It thus appears that the P300 correlated with visual awareness in this experiment.

**Figure 3.**
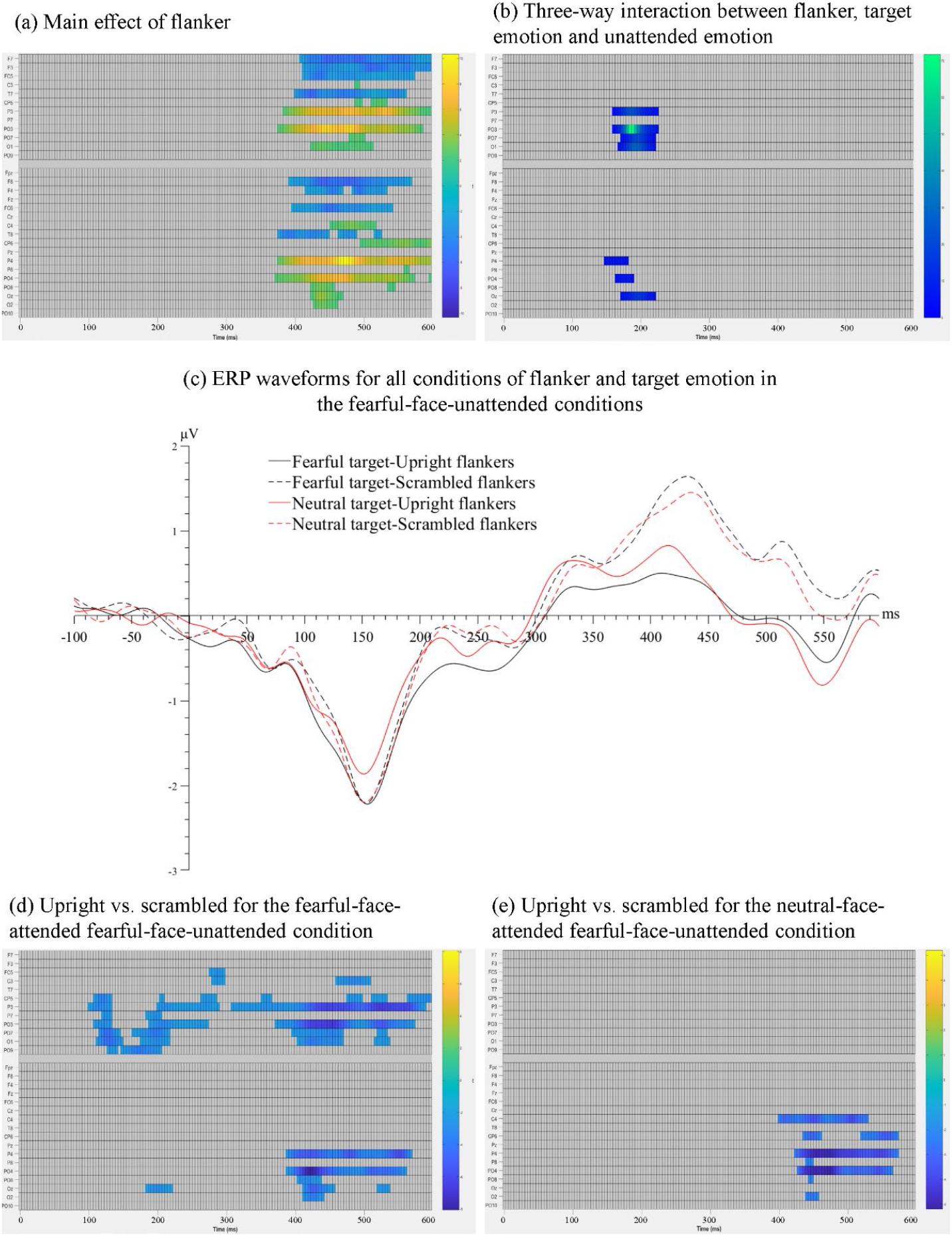
Raster plots for (a) the significant main effect of flanker and (b) the significant three-way interaction between target emotion, unattended emotion and flanker. (c) ERP waveforms of the four conditions of target emotion and flanker, when the unattended face was a fearful face, over the average of electrodes P3/4 and PO3/4, the two pairs of electrodes showing the maximal flanker effects. Raster plots for the pairwise comparisons between the upright flanker condition and the scrambled flanker condition (d) when the target face was a fearful face and (e) when the target face was a neutral face, in trials where the unattended face was a fearful face. Data in trials where the unattended face was a neutral face were not shown.

To follow up the significant three-way interaction, we ran 2(target emotion) x 2(flanker) ANOVAs separately for the condition where the unattended face was a fearful face and for the condition where it was a neutral face. Results showed that the interaction between target emotion and flanker was significant only when the unattended face expressed fear, *Fs* > 4.39, *ps* < .035. However, this two-way interaction was not significant when the unattended face was a neutral face, *ps* > .308. Thus, we then compared the upright trials to the scrambled trials separately for two different target emotions (fearful and neutral) in conditions where the unattended face was a fearful face.

When the target face was a fearful face, there were two negative clusters when contrasting upright against scrambled conditions: an earlier negative cluster (CP5, P3, P7, PO3, PO7, PO9, O1, Oz) spanning from 98 to 286ms (spatial peak: PO7, temporal peak: 126ms), *ts* > 2.09, *ps* < .029, and a later negative cluster (C3, CP5, P3/4, PO3/4, PO7/8, O1/2, Oz) spanning from 306 to 594ms (spatial peak: PO7, temporal peak: 422ms), *ts* > 2.13, *ps* < .002, see Figure 3d. By contrast, when the target face was a neutral face, we only found one significant negative cluster in the upright-scrambled comparison, which spanned from 398-570ms over C4, CP6, P4, PO4 and O2 (spatial peak: PO4, temporal peak: 470ms), *ts* > 2.12, *ps* < .011, see Figure 3e. That is, the late flanker effect on the P300 (i.e., a positivity for the scrambled flanker condition) was found regardless of target emotion, which is likely an effect related to awareness of the target face, as noted above. However, when the attended target face was a fearful face, there was an additional earlier negativity for the upright flanker condition, compared to the scrambled flanker condition, and this early effect was not found when the target face was a neutral face.

No other omnibus effect was significant, *ps* > .167.

## Discussion

In the current study, we examined the processing of fearful and neutral faces using a visual crowding paradigm. Upright angry face flankers and scrambled face flankers were used to manipulate awareness. Our results provide evidence that the P300 over parietal electrodes may index awareness of the central target faces in this paradigm. Additionally, there seemed to be a congruency effect related to negative emotions in an earlier time window (∼120ms) over posterior electrodes. Interestingly, this effect was observed only when the central face in the unattended clutter was a fearful face and not when it was a neutral face.

In our experiment, we intended to manipulate visual awareness of central crowded faces by modifying flankers whereby upright flankers would reduce visual awareness of central faces, compared to other conditions. Previous studies have shown that when the similarity between the target and flankers was high, for example when they were both upright human face images, the crowding effect was stronger for the central face (e.g., [13,15]). In support of this, we found that participants were more likely to report seeing the central target face in the scrambled relative to the upright flanker condition. Importantly, this behavioural finding was accompanied by an enhanced electrophysiological activity in the P300 time window for the scrambled flanker condition compared to the upright flanker condition.

Previous research has shown that the P300 is associated with a higher-level, reportable form of awareness, and it is often enhanced in situations where participants reported seeing a stimulus, compared to when they missed it [16,29]. By contrast, the earlier VAN has been suggested to index awareness associated with the mere perceptual experience of stimuli, and the reporting of the stimuli is not a requirement for this component [16]. From this we infer that the current emotion judgement task may require a rather effortful evaluation of the target stimuli. Consequently, awareness may occur only during a later stage of processing (i.e., the P300 time range) where such an evaluative appraisal of the stimuli took place.

With an onset of ∼300ms post-stimulus, the P300 is also characterised by a variety of post-perceptual processes [17,30,31]. For example, a previous study found that the P300 was enhanced by task-relevant targets but it was completely absent for task-irrelevant stimuli, even though participants were aware of them [30]. In this case, the P300 appeared to be a correlate for task relevancy of the stimuli, rather than awareness *per se*. Similarly, in a change blindness study by Scrivener et al. [32], the P300 was found to be strengthened when changes were localised, compared to when they were merely detected, whereas the earlier VAN was found to be comparable between a successful detection and a correct localisation of the changes. It was concluded that the VAN was linked to the perceptual and an early stage of awareness whereas the P300 was indicative of higher level of task processing [32]. Therefore, while the enhanced P300 in our scrambled flanker condition could be associated with awareness, it may also reflect post-perceptual processes that were not directly related to awareness.

Contrary to predictions, we did not find an earlier enhanced negativity (the VAN) for the scrambled condition. One possible explanation for this finding is that any enhanced negativity related to awareness in the scrambled condition may have been offset by the decrease in the signals due to the smaller number of actual face images. That is, the presence of six additional faces (flankers) in the upright condition may have increased the neural activity in the N2 time range, compared to the scrambled flanker condition where only one face was presented. Consequently, no significant increase or decrease in the ERPs could be found in this time window when comparing the two flanker conditions. One may contend that using upright faces as flankers can pose an issue for interpretation but we deemed their use necessary in the condition intended to impede awareness the most. Future studies are needed to test whether these results hold true for non-face visual stimuli.

Interestingly, at around 120ms, a fearful target face enhanced the ERPs in the upright flanker condition compared to the scrambled flanker condition. This enhancement was not found for a neutral target face. That is, signals in this early time window were stronger when both the target face and the flanker faces expressed negative emotions (fear and anger, respectively). However, when the target face was a neutral face, there was no difference between the two flanker conditions. Thus, there seems to be a congruency effect between the central target and the flankers, which, importantly, occurred only for negative emotions.

Because we only used angry faces as a type of upright face flankers, there was only one possible congruent combination of the target and flankers (i.e., fearful face target and angry face flankers), we cannot be sure that such a congruency effect was specific to negative emotions. Further studies are needed to test whether this congruency effect can be found for other emotions (e.g., positive emotions). Nevertheless, it seems that in a visual crowding paradigm, the early neural processing of the crowded target can be affected by its relation with the flankers, or particularly the similarity between them. Any modulations related to awareness induced by the upright flankers may have been too small, if not completely absent, during the early processing of the stimuli in this study.

Finally, the above interaction between flankers and target emotion was found only when the unattended face was a fearful face and not when it was a neutral face. This is consistent with claims that unattended fearful expressions can facilitate the processing of the attended information [33]. A series of studies conducted by Bertini and colleagues (for reviews see [33,34]) on hemianopic patients showed that, task-irrelevant emotional faces presented in the blind visual field of the patients can be processed nonconsciously.

Specifically, some patients who were blind in one half of the visual field demonstrated better discrimination of visual stimuli presented in their intact visual field, when a fearful face, compared to a neutral or a happy face, was presented simultaneously in the blind field [34]. It was argued that unattended and unseen fearful faces were processed via alternate neural pathways and subsequently facilitated the processing of attended stimuli [35]. Our current findings are consistent with this claim by showing that, only when the unattended face was a fearful face, participants’ neural activity could distinguish between a fearful and a neutral target face, even in situations where the perceivability of the target face was very low (i.e., in the upright flanker condition).

In conclusion, during visual crowding for faces, early neural responses at about 120ms post-stimulus seem to be affected by the perceptual characteristics of the stimuli, specifically the target-flanker congruency, whereas neural activity at a later stage is associated with post-perceptual evaluation of the crowded stimulus.

## Acknowledgement

Zeguo Qiu and Benjamin J. Muehlebach were supported by Graduate Scholarships provided by University of Queensland.

## CRediT authorship contribution statement

**Zeguo Qiu**: Conceptualization, Methodology, Formal analysis, Investigation, Data curation, Writing - Original Draft, Writing – Review & Editing, Visualization, Project administration. **Dihua Wu**: Investigation. **Benjamin J. Muehlebach**: Writing - Review & Editing.

## Declaration of generative AI in scientific writing

The authors declare that no generative artificial intelligence (AI) and AI-assisted technologies were used in the writing process.

## Data availability

Data will be made available on request.

## References

[1] R. J. Compton, 2003. The interface between emotion and attention: A review of evidence from psychology and neuroscience. Behavioral and cognitive neuroscience reviews, 2(2), 115–129. https://doi.org/10.1177/1534582303002002003

[2] S. Schindler, F. Bublatzky, 2020. Attention and emotion: An integrative review of emotional face processing as a function of attention. Cortex, 130, 362–386. https://doi.org/10.1016/j.cortex.2020.06.010

[3] M. Eimer, M. Kiss, 2007. Attentional capture by task-irrelevant fearful faces is revealed by the N2pc component. Biological psychology, 74(1), 108–112. https://doi.org/10.1016/j.biopsycho.2006.06.008

[4] Z. Qiu, S. I. Becker, A. J. Pegna, 2022. The Effects of Spatial Attention Focus and Visual Awareness on the Processing of Fearful Faces: An ERP Study. Brain Sciences, 12(7), 823. https://doi.org/10.3390/brainsci12070823

[5] G. Stefanics, G. Csukly, S. Komlósi, P. Czobor, I. Czigler, 2012. Processing of unattended facial emotions: a visual mismatch negativity study. Neuroimage, 59(3), 3042–3049. https://doi.org/10.1016/j.neuroimage.2011.10.041

[6] Z. Qiu, S. I. Becker, A. J. Pegna, 2022. Spatial attention shifting to emotional faces is contingent on awareness and task relevancy. Cortex, 151, 30–48. https://doi.org/10.1016/j.cortex.2022.02.009

[7] A. J. Pegna, A. Darque, C. Berrut, A. Khateb, 2011. Early ERP modulation for task-irrelevant subliminal faces. Frontiers in psychology, 2, 88. https://doi.org/10.3389/fpsyg.2011.00088

[8] Z. Qiu, J. Zhang, A. J. Pegna, 2023. Neural processing of lateralised task-irrelevant fearful faces under different awareness conditions. Consciousness and Cognition, 107, 103449. https://doi.org/10.1016/j.concog.2022.103449

[9] D. Baier, M. Kempkes, T. Ditye, U. Ansorge, 2022. Do Subliminal Fearful Facial Expressions Capture Attention?. Frontiers in Psychology, 1782. https://doi.org/10.3389/fpsyg.2022.840746

[10] Z. Qiu, J. Jiang, S. I. Becker, A. J. Pegna, 2023. Attentional capture by fearful faces requires consciousness and is modulated by task-relevancy: A dot-probe EEG study. Frontiers in Neuroscience, 17. https://doi.org/10.3389/fnins.2023.1152220

[11] E. Tipura, A. J. Pegna, 2022. Subliminal emotional faces do not capture attention under high attentional load in a randomized trial presentation. Visual Cognition, 30(4), 280–288. https://doi.org/10.1080/13506285.2022.2060397

[12] D. Whitney, D. M. Levi, 2011. Visual crowding: A fundamental limit on conscious perception and object recognition. Trends in cognitive sciences, 15(4), 160–168. https://doi.org/10.1016/j.tics.2011.02.005

[13] A. V. Kalpadakis-Smith, V. Goffaux, J. A. Greenwood, 2018. Crowding for faces is determined by visual (not holistic) similarity: Evidence from judgements of eye position. Scientific reports, 8(1), 1–14. https://doi.org/10.1038/s41598-018-30900-0

[14] M. Gong, L. J. Smart, 2021. The anger superiority effect revisited: a visual crowding task. Cognition and Emotion, 35(2), 214–224. https://doi.org/10.1080/02699931.2020.1818552

[15] E. G. Louie, D. W. Bressler, D. Whitney, 2007. Holistic crowding: Selective interference between configural representations of faces in crowded scenes. Journal of Vision, 7(2), 24–24. https://doi.org/10.1167/7.2.24

[16] J. Förster, M. Koivisto, A. Revonsuo, 2020. ERP and MEG correlates of visual consciousness: The second decade. Consciousness and cognition, 80, 102917. https://doi.org/10.1016/j.concog.2020.102917

[17] M. A. Cohen, K. Ortego, A. Kyroudis, M. Pitts, 2020. Distinguishing the neural correlates of perceptual awareness and postperceptual processing. Journal of Neuroscience, 40(25), 4925–4935. https://doi.org/10.1523/JNEUROSCI.0120-20.2020

[18] M. Del Zotto, A. J. Pegna, 2015. Processing of masked and unmasked emotional faces under different attentional conditions: an electrophysiological investigation. Frontiers in psychology, 6, 1691. https://doi.org/10.3389/fpsyg.2015.01691

[19] N. Faivre, V. Berthet, S. Kouider, 2012. Nonconscious influences from emotional faces: A comparison of visual crowding, masking, and continuous flash suppression. Frontiers in psychology, 3, 129. https://doi.org/10.3389/fpsyg.2012.00129

[20] H. M. Sun, B. Balas, 2015. Face features and face configurations both contribute to visual crowding. Attention, Perception, & Psychophysics, 77, 508–519. https://doi.org/10.3758/s13414-014-0786-0

[21] C. Peng, C. Hu, Y. Chen, 2018. The temporal dynamic relationship between attention and crowding: Electrophysiological evidence from an event-related potential study. Frontiers in Neuroscience, 12, 844. https://doi.org/10.3389/fnins.2018.00844

[22] J. Peirce, J. R. Gray, S. Simpson, M. MacAskill, R. Höchenberger, H. Sogo,… J. K. Lindeløv, 2019. PsychoPy2: Experiments in behavior made easy. Behavior research methods, 51, 195–203. https://doi.org/10.3758/s13428-018-01193-y

[23] O. Langner, R. Dotsch, G. Bijlstra, D. H. Wigboldus, S. T. Hawk, A. D. Van Knippenberg, 2010. Presentation and validation of the Radboud Faces Database. Cognition and emotion, 24(8), 1377–1388. https://doi.org/10.1080/02699930903485076

[24] J. A. Greenwood, M. Szinte, B. Sayim, P. Cavanagh, 2017. Variations in crowding, saccadic precision, and spatial localization reveal the shared topology of spatial vision. Proceedings of the National Academy of Sciences, 114(17), E3573–E3582. https://doi.org/10.1073/pnas.1615504114

[25] A. Delorme, S. Makeig, 2004. EEGLAB: an open source toolbox for analysis of single-trial EEG dynamics including independent component analysis. Journal of neuroscience methods, 134(1), 9–21. https://doi.org/10.1016/j.jneumeth.2003.10.009

[26] J. Lopez-Calderon, S. J. Luck, 2014. ERPLAB: an open-source toolbox for the analysis of event-related potentials. Frontiers in human neuroscience, 8, 213. https://doi.org/10.3389/fnhum.2014.00213

[27] E. C. Fields, G. R. Kuperberg, 2020. Having your cake and eating it too: Flexibility and power with mass univariate statistics for ERP data. Psychophysiology, 57(2), e13468. https://doi.org/10.1111/psyp.13468

[28] D. M. Groppe, T. P. Urbach, M. Kutas, 2011. Mass univariate analysis of event-related brain potentials/fields II: Simulation studies. Psychophysiology, 48(12), 1726–1737. https://doi.org/10.1111/j.1469-8986.2011.01272.x

[29] C. Dembski, C. Koch, M. Pitts, 2021. Perceptual awareness negativity: a physiological correlate of sensory consciousness. Trends in Cognitive Sciences, 25(8), 660–670. https://doi.org/10.1016/j.tics.2021.05.009

[30] M. A. Pitts, J. Padwal, D. Fennelly, A. Martínez, S. A. Hillyard, 2014. Gamma band activity and the P3 reflect post-perceptual processes, not visual awareness. Neuroimage, 101, 337–350. https://doi.org/10.1016/j.neuroimage.2014.07.024

[31] R. Rutiku, M. Martin, T. Bachmann, J. Aru, 2015. Does the P300 reflect conscious perception or its consequences?. Neuroscience, 298, 180–189. https://doi.org/10.1016/j.neuroscience.2015.04.029

[32] C. L. Scrivener, A. Malik, J. Marsh, M. Lindner, E. B. Roesch, 2019. An EEG study of detection without localisation in change blindness. Experimental brain research, 237, 2535–2547. https://doi.org/10.1007/s00221-019-05602-2

[33] C. Bertini, E. Làdavas, 2021. Fear-related signals are prioritised in visual, somatosensory and spatial systems. Neuropsychologia, 150, 107698. https://doi.org/10.1016/j.neuropsychologia.2020.107698

[34] E. Làdavas, C. Bertini, 2021. Right hemisphere dominance for unconscious emotionally salient stimuli. Brain Sciences, 11(7), 823. https://doi.org/10.3390/brainsci11070823

[35] C. Bertini, R. Cecere, E. Làdavas, 2013. I am blind, but I “see” fear. Cortex, 49(4), 985–993. https://doi.org/10.1016/j.cortex.2012.02.006

